# Cystin gene mutations cause autosomal recessive polycystic kidney disease associated with altered *Myc* expression

**DOI:** 10.1101/2020.02.18.946285

**Authors:** Chaozhe Yang, Amber K. O’Connor, Robert A. Kesterson, Jacob A. Watts, Amar J. Majmundar, Daniela A. Braun, Monkol Lek, Kristen M. Laricchia, Hanan M. Fathy, Shirlee Shril, Friedhelm Hildebrandt, Lisa M. Guay-Woodford

## Abstract

Mutation of the *Cys1* gene underlies the renal cystic disease in the *Cys1*^*cpk*/*cpk*^ (*cpk)* mouse that phenocopies human autosomal recessive polycystic kidney disease (ARPKD). Cystin, the protein product of *Cys1*, is expressed in the primary apical cilia of renal ductal epithelial cells. In previous studies, we showed that cystin regulates *Myc* expression via interaction with the tumor suppressor, necdin. Here, we demonstrate rescue of the *cpk* renal phenotype by kidney-specific expression of a cystin-GFP fusion protein encoded by a transgene integrated into the *Rosa26* locus. In addition, we show that expression of the cystin-GFP fusion protein in collecting duct cells down-regulates expression of *Myc* in *cpk* kidneys. Finally, we report the first human patient with an ARPKD phenotype due to homozygosity for a predicted deleterious splicing defect in *CYS1*. These findings suggest that mutations in the *Cys1* mouse and *CYS1* human orthologues cause an ARPKD phenotype that is driven by overexpression of the *Myc* proto-oncogene.

**Translational Statement:** The cystin-deficient *cpk* mouse is a model for the study of autosomal recessive polycystic kidney disease (ARPKD). We show that the *cpk* mouse phenotype is associated with altered *Myc* expression. To date, the clinical relevance of cystin deficiency to human disease was unclear, due to the absence of ARPKD cases associated with *CYS1* mutations. We report the first case of ARPKD linked to a *CYS1* mutation disrupting normal splicing. These findings confirm the relevance of cystin deficiency to human ARPKD, implicate *Myc* in disease initiation or progression, and validate the *cpk* mouse as a translationally relevant disease model.

## Introduction

Autosomal recessive polycystic kidney disease (ARPKD; MIM 263200) affects 1:26,500 live births.^1^ Cohort studies indicate that 75-80% of patients with typical ARPKD have mutations in the Polycystic Kidney and Hepatic Disease 1 (*PKHD1*) gene.^2–5^ Mutations in the *DZIP1L* gene account for less than 0.1% of affected patients,^6^ while mutations in other hepato-renal fibrocystic disease genes, eg. *HNF1B*, *PKD1*, *NPHP2*, *NPHP3*, and *NPHP13*, can phenocopy ARPKD.^7^ For reasons yet to be explained, mice with targeted disruption of *Pkhd1* exhibit little or no kidney disease.^8–15^ In the absence of a *Pkhd1* mutant mouse model that accurately recapitulates the human disease phenotype, the *cpk* mouse carrying a spontaneous truncating mutation in *Cys1* has been the most widely studied mouse model of ARPKD.^16,17^ Cystin, the *Cys1* gene product, is a 145-amino acid cilia-associated protein that is expressed in mouse embryonic kidney and liver ductal epithelium.^18^ Disruption of cystin function results in elevated *Myc* expression in collecting duct epithelial cells^19–22^ and increased cell proliferation.^19,23^ In previous work, we have demonstrated that in renal collecting duct epithelia, cystin physically interacts with necdin in a regulatory complex that modulates *Myc* expression.^24^

Cystin deficiency-associated disruption of ciliary signaling and/or overexpression of *Myc* is associated with aberrant SMAD3 phosphorylation,^25^ overexpression of *Fos* and *Kras* proto-oncogenes,^19–21^ elevated levels of growth factors,^26^ aberrant localization and abundance of the epidermal growth factor receptor (EGFR) on the apical surface of collecting duct cells^27^, abnormal levels of basement membrane components,^28–30^ and epithelial cell adhesion molecules.^31,32^ Until now, the relevance of these effects of cystin deficiency for human disease was unclear in the absence of ARPKD patients with mutations in human *CYS*. Here we present the first case of human ARPKD due to homozygosity for a *CYS1* mutation, in this case predicted to cause defective splicing. We also show that complementation of defective *Cys1* in the kidneys of *Cys1*^*cpk/cpk*^ (*cpk*) mice rescues both *Myc* overexpression and the collecting duct cyst phenotype. These studies suggest that up-regulation of *Myc* expression *in vivo* may play a central role in the pathogenesis of mouse recessive polycystic kidney disease, with important implications for human ARPKD.

## Results

### Phenotypic rescue of cpk mice by kidney-specific expression of a cystin-GFP fusion protein

We generated a conditional expression *Cys1* transgenic *Cys1^cpk/cpk^ (cpk)* mouse line carrying a *Cys1-GFP* transgene knock-in at the *Rosa26* locus. In these mice, *Cys1-GFP* transgene expression is precluded by the presence of a loxP-flanked termination sequence consisting of a PGK-Neo cassette (Figure1A, T^OFF^ allele). The *Cys1*-GFP transgene is expressed by the ROSA26 promoter only after *Cre*-mediated deletion of the loxP-flanked PGK-Neo cassette (Figure1A, T^ON^ allele). We crossed *Rosa26-Cys1-GFP* mice with *Cys1*^*cpk*/+^ mice to generate *Cys1*^*cpk*/+^;*Rosa26-Cys1-GFP* mice, which were then crossed with *Ksp-Cre* transgenic mice^33^ to generate *Cys1*^*cpk*/+^;*Rosa26-Cys1-GFP;Ksp-Cre* progeny. In these mice, *Cre* expression, controlled by the *Ksp*-cadherin regulatory elements, occurs exclusively in the developing distal renal tubular epithelium and the genitourinary tract^34^ resulting in high level expression of the cystin-GFP fusion protein in the collecting ducts and loops of Henle and low or no expression in the proximal tubules. Finally, the rescue experiments were carried out by crossing *Cys1*^*cpk*/+^;*Rosa26-Cys1-GFP;Ksp-Cre* mice with *Cys1*^*cpk*/+^ mice. Genotype-confirmed *Cys1^cpk/cpk^;Rosa26-Cys1-GFP;Ksp-Cr*e experimental “rescue” (R) mice were compared to their *Cys1^+/+^;Rosa26-Cys1-GFP;Ksp-Cre* control (C) littermates (Figure 1B). While *cpk* mice are characteristically smaller than wild-type littermates and die by 21 days of age,^35^ no differences in body size or survival were observed between the rescue (R) mice and their littermate controls (Figure 1C, left panel). Kidney sizes at postnatal days 14 and 21 were not significantly different in R and wild-type (WT) mice (Figure 1C, right panel), while age-matched *Cys1*^*cpk*/*cpk*^ (*cpk*) mice exhibited the characteristic cystic kidney phenotype. These results indicate that the gross renal phenotype of the *cpk* mouse was rescued by kidney-specific expression of cystin-GFP.

**Figure 1.**
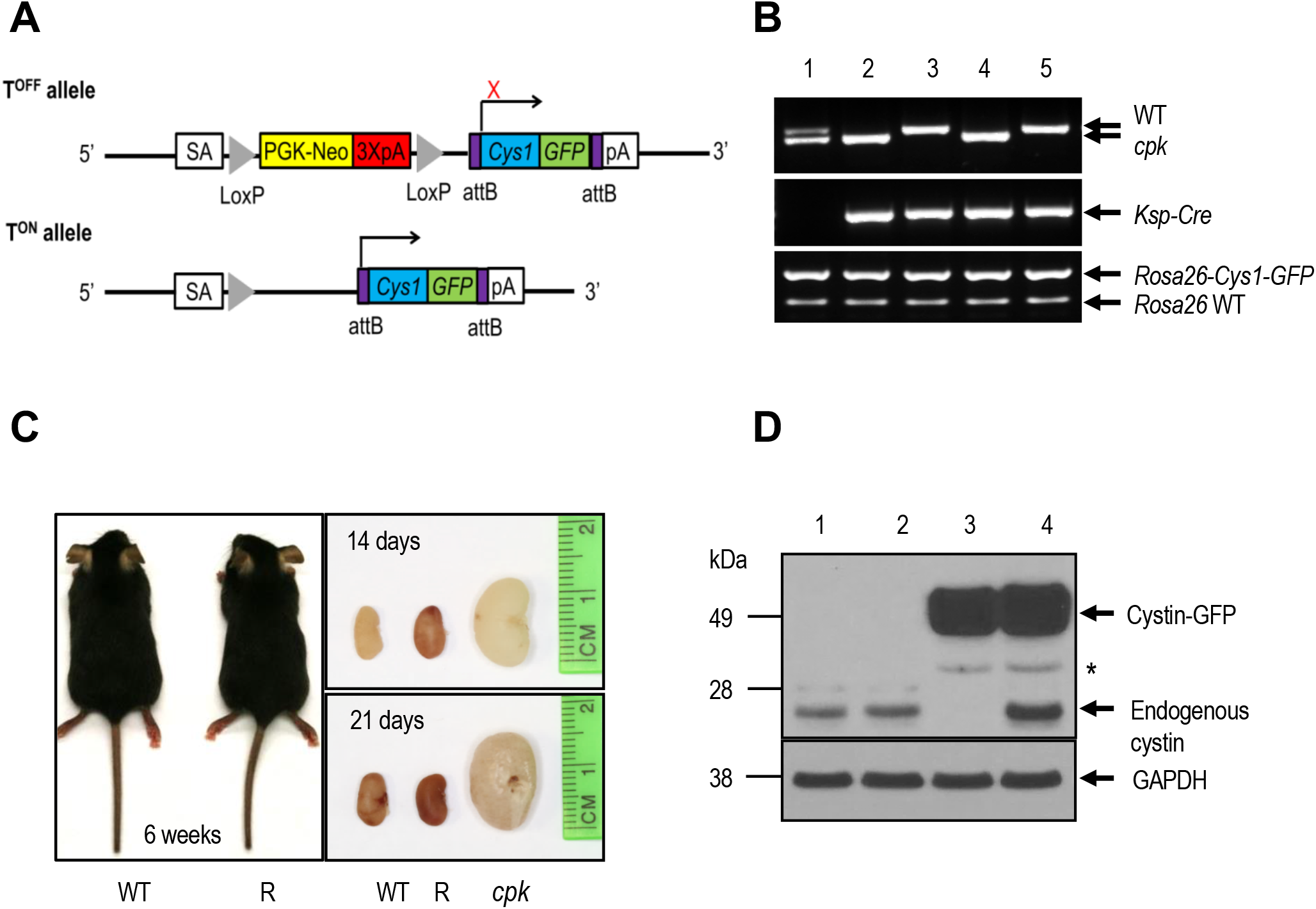
(**A**) Schematic diagram showing the *Cys1-GFP* transgene knock-in at the *Rosa26* locus (*Rosa26-Cys1-GFP* allele) before (T^OFF^) and after (T^ON^) deletion of a PGK Neo cassette (yellow rectangle) flanked by LoxP sites (gray triangles). In the T^OFF^ configuration, expression of *Cys1-GFP* is prevented by the PGK Neo cassette. In cells expressing a *Ksp-Cre* transgene, Cre-mediated recombination deletes PGK Neo and *Cys1-GFP* is expressed (T^ON^). SA: splice acceptor, PKG-Neo: Phosphoglycerate kinase promoter driving a neomycin resistance gene followed by 3 polyA signals (3XpA, red rectangle). The purple boxes flanking *Cys1-GFP* are attB sites. (**B)** PCR-based genotyping of *Rosa26-Cys1-GFP*, *Ksp-Cre* and *Cys1* alleles: lane 1 – *Cys1^cpk/+^;Rosa26-Cys1-GFP/+* mice; lane 2 and 4 – *Cys1^cpk/cpk^;Rosa26-Cys1-GFP; Ksp-Cre/+* mice; lanes 3 and 5 – *Cys^+/+^;Rosa26-Cys1-GFP;Ksp-Cre/+* mice. (**C**) *Cys1-GFP* rescue of gross phenotypes in *cpk* mice. Six week-old wild-type (WT) and *Cys1^cpk/cpk^;Rosa26-Cys1-GFP; Ksp-Cre/+* rescued (R) mice are of equivalent size. Examination of kidneys from WT, R and *cpk* mice at 14 and 21 days of age show equivalent sizes of R and WT kidneys, with both markedly smaller than *cpk* kidneys. (**D**) Western blot analysis of total kidney protein from 6 week-old mice: lane 1 – C57Bl/6 wild-type mice; lane 2 – *Cys^cpk/+^*;*Rosa26-Cys1-GFP* mice; lane 3 – R mice; lane 4 – C mice. Mouse cystin is 145 amino acids long but migrates aberrantly at ~25 kDa on SDS-PAGE. Cystin-GFP (arrow, ~50 kDa) and endogenous cystin (arrowhead, ~25 kDa) were detected using polyclonal rabbit anti-cystin antibody. GAPDH served as an internal protein loading and transfer control. The asterisk indicates non-specific bands.

### Expression of cystin-GFP fusion protein in the kidneys of rescued cpk mice

We examined the expression of the cystin-GFP fusion protein in the kidneys of R mice. Endogenous cystin was detectable in the kidneys of both WT and C mice and absent from the kidneys of R mice (Figure 1D). The cystin-GFP fusion protein of ~50kDa was detected in R and C mice (Figure 1D, lanes 3 and 4). These results demonstrate that cystin-GFP expression was associated with Cre-mediated excision of the PGK-Neo cassette.

Dual immunofluorescence staining with antibodies against GFP and aquaporin-2 (AQP2) was used to examine cystin-GFP expression in nephron segments of kidneys from R and C mice (Figure 2). AQP2 is expressed primarily on apical cell membranes of collecting duct cells.^36,37^ The cystin-GFP fusion protein was detected in AQP2-positive collecting ducts of R mice (Figure 2C and I) and C mice (Figure 2B and H), while cystin-GFP was absent in the *Rosa26-Cys1-GFP* mice that do not carry *Ksp-Cre* transgene (Figure 2A and G). Co-localization of AQP2 and cystin-GFP demonstrated cystin-GFP fusion protein expression in the collecting duct cells.

**Figure 2.**
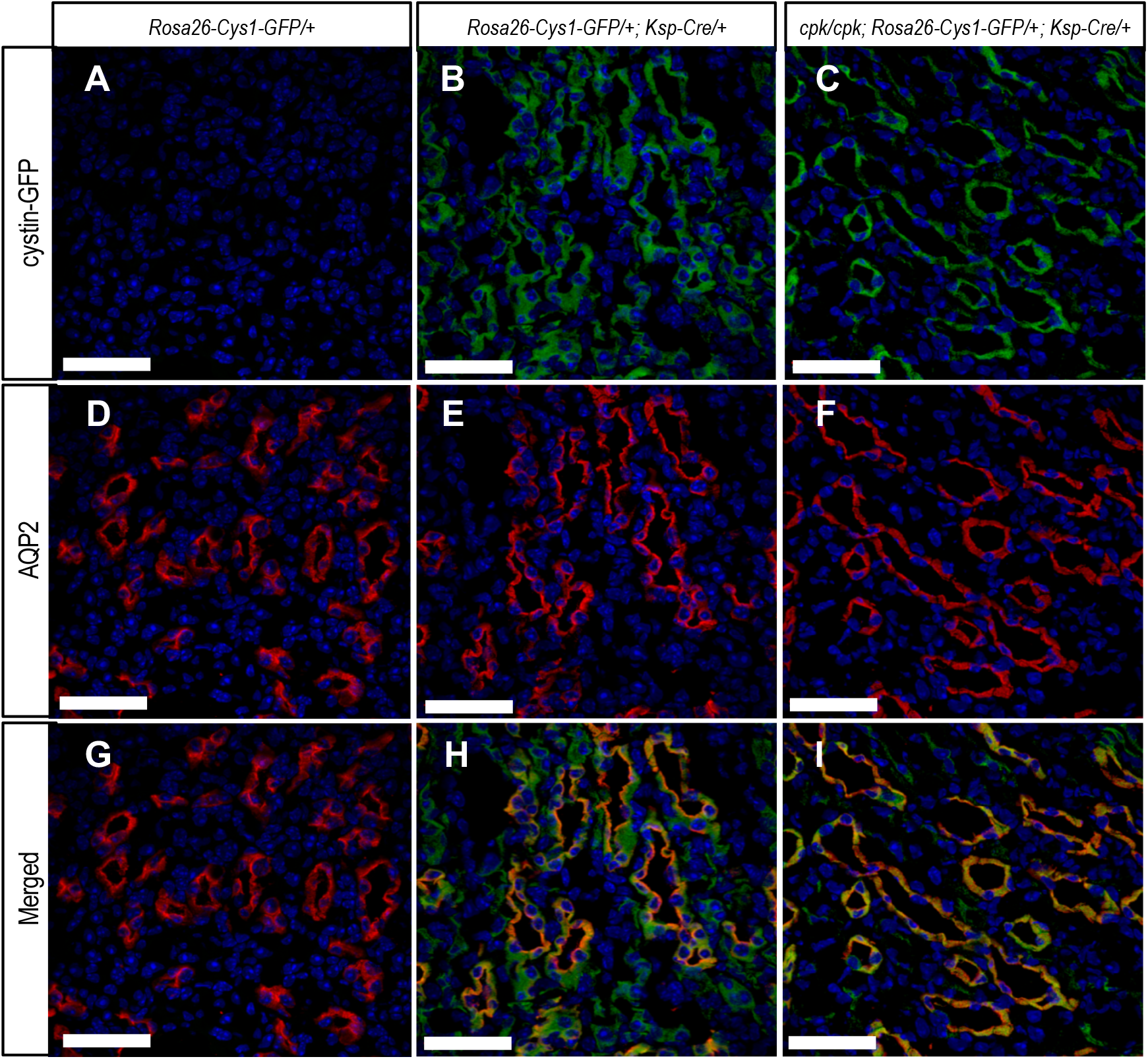
Immunohistochemical detection of cystin-GFP fusion protein (green, **A-C**), AQP2 (red, **D-F**) and merged (**G-I**) in kidney tissues from 6 week-old mice of the indicated genotypes. The *Rosa26-Cys1-GFP/+* mice (**A,D,G**) do not express cystin-GFP fusion protein due to the absence of the *Ksp-Cre* transgene. The *Rosa26-Cys1-GFP/+; Ksp-Cre/+* mice (**B,E,H**) have a wild-type Cys1 gene and express cystin-GFP fusion protein in Cre-positive cells. The *cpk/cpk; Rosa26-Cys1-GFP/+;Ksp-Cre/+* rescue mice (**C,F,I**) express cystin-GFP fusion protein in Cre-positive cells. Cell nuclei are stained with DAPI (blue). Scale bars equal 10 μm. Images are representative of tissue sections from 3 animals.

### Histological evaluation of cystogenesis in rescued cpk mice

The gross evaluation of kidneys from R mice suggested that the renal histology would be normal. Histological evaluation showed that, while the majority of the nephrons in R kidneys appeared to have normal dimensions, occasional cystic structures were present (Figure 3C and F). Using DBA and LTA lectins, markers of distal and proximal tubules, respectively,^38,39^ we observed that the cystic structures stained with LTA (Figure 3I). These findings suggest that while expression of the cystin-GFP in collecting ducts markedly attenuated the *cpk* renal phenotype, sporadic cyst formation did occur in proximal tubular segments of these kidneys (Figure 3I).

**Figure 3.**
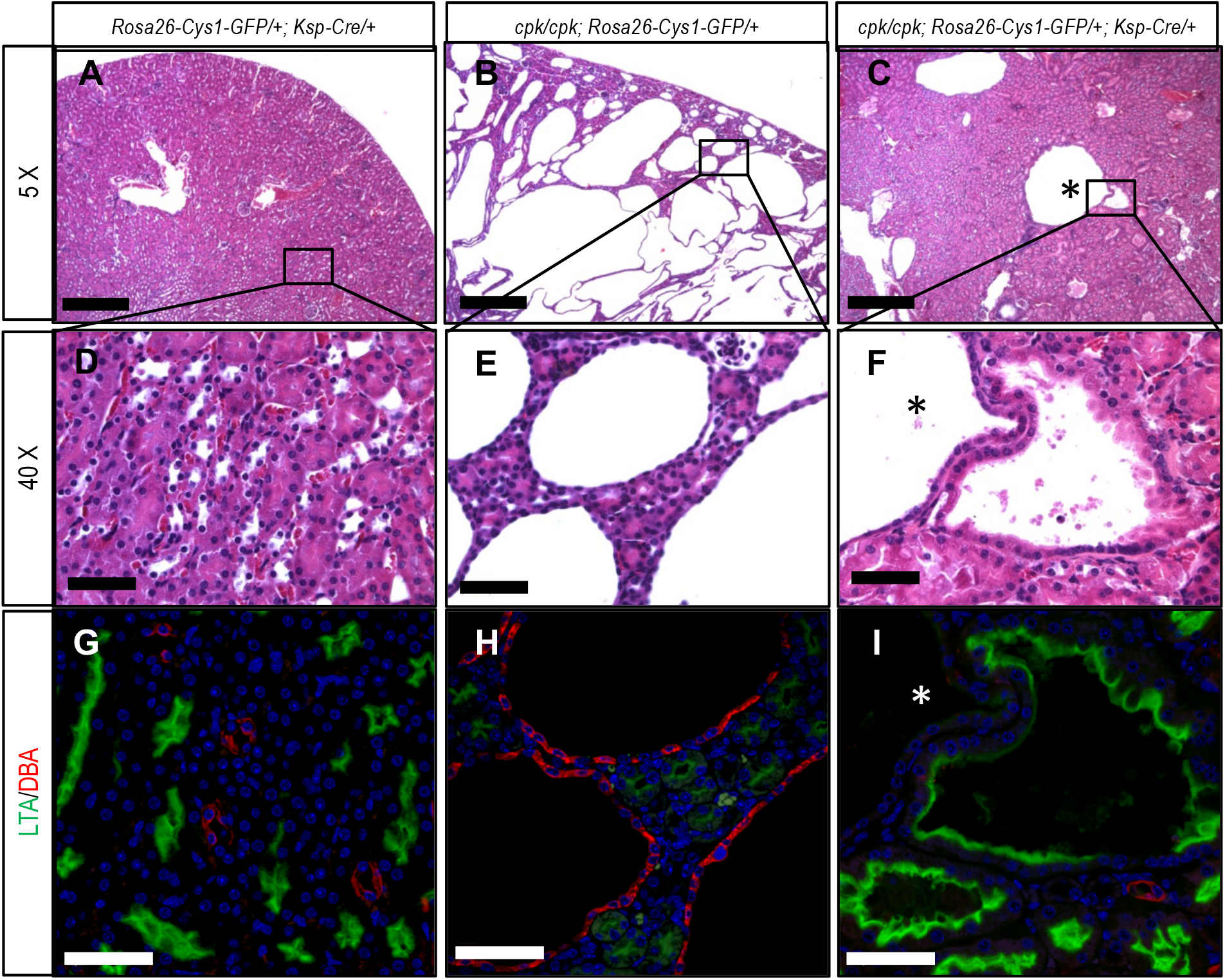
Kidney histology and differential lectin staining. Formalin fixed, paraffin embedded 5 μm sections of kidney tissues from 6 week-old mice of the indicated genotypes were stained with H&E and examined by light microscopy (**A-F**) or stained with lectins LTA (green, proximal tubules) and DBA (red, distal tubules) and examined by immunofluorescence microscopy (**G,H,I**). Panels A-C are 5X magnification with scale bars equal to 100 μm. Boxed areas are shown at 40X magnification in corresponding panels D-F with scale bars equal to 20 μm. Lectin staining was performed on serial sections corresponding to H&E stained samples. Cell nuclei were stained with DAPI. Scale bars in panels G-I are equal to 10 μm. The asterisk (*) identifies a large cyst that was not stained by either LTA or DBA. Images are representative of tissue sections from 3 animals.

### Expression of Myc in rescued cpk mice

*Myc* overexpression in *cpk* kidneys is well-documented.^19–22^ In previous work we demonstrated that cystin physically interacts with the DNA-binding protein necdin in a regulatory complex that binds to the *Myc* P1 promoter.^24^ Necdin enhances *Myc* promoter activity and cystin antagonizes this effect. In a previous report, we proposed that *Myc* up-regulation in *cpk* kidneys results directly from disruption of the cystin-necdin interaction. In the current study, we examined the relative abundance of C-MYC protein in the kidneys of 14-day old R mice as compared to *cpk* mice. Quantitative immunoblotting revealed comparable levels of C-MYC in WT and R kidneys that were markedly lower than in kidneys of *cpk* mice (Figure 4). These results demonstrate that transgene rescue of the cystic kidney phenotype in *cpk* mice is associated with down-regulation of C-MYC protein expression, suggesting a central role for *Myc* overexpression in the renal cystogenesis in this mouse model.

**Figure 4.**
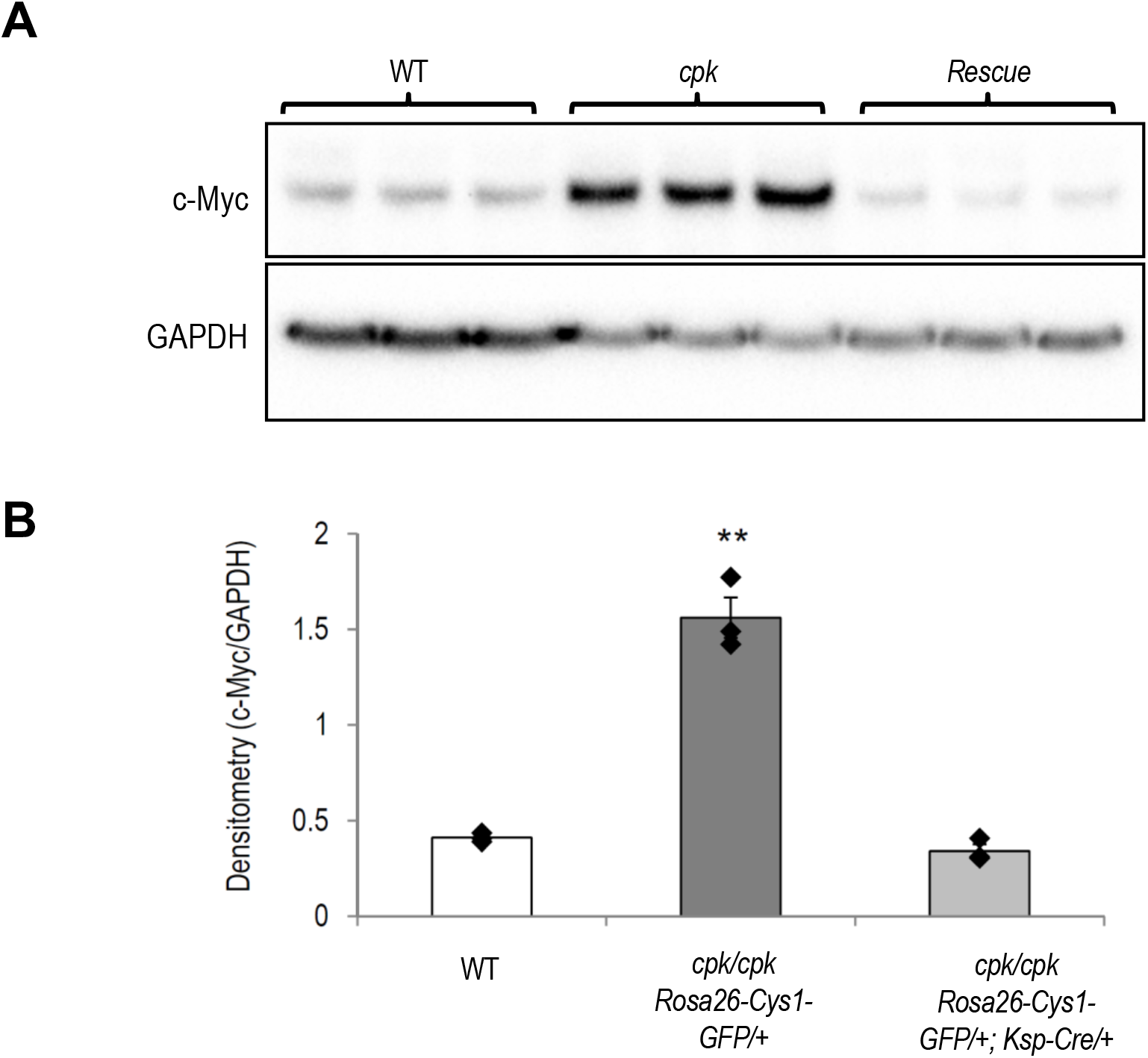
Immunoblot analysis of C-MYC protein expression in kidneys from wild-type (WT), *cpk/cpk*;*Rosa26-Cys1-GFP/+* (*cpk*) and *cpk/cpk*;*Rosa26-Cys1-GFP/+;Ksp-Cre/+* (Rescue) mice. (**A**) Total kidney lysates were immunoblotted using anti-C-MYC and anti-GAPDH antibodies. (**B**) C-MYC band intensity was normalized with GAPDH. Data represents mean ± S.E; Y-axis values indicate C-MYC/GAPDH band intensity ratio; **P< 0.01, vs. others, ANOVA, n = 3 each group.

### A case of human ARPKD associated with homozygous CYS mutation

While the *cpk* mouse phenotype recapitulates important clinical features of ARPKD, to date no human cases of the disease have been linked to mutation of the *CYS1* gene. Recently, a 5-year-old male patient (subject B783), the offspring of consanguineous parents, was evaluated for monogenic cystic renal disease. The patient presented with polyuria and polydipsia and ultrasound examination revealed multiple medullary and cortical cysts consistent with polycystic kidney disease. Renal function was mildly reduced (creatinine 0.6 mg/dL) for a child of the subject’s age. DNA samples obtained from from the proband and both parents were analyzed by trio whole exome sequencing (TRIO-WES). Homozygosity mapping of exome variant data for B783 indicated 108 Mbp of homozygosity by descent (Figure 5A), suggesting that the parents are approximately fourth degree relatives. Based on this mapping, it was hypothesized that a biallelic gene mutation residing within a homozygous peak region caused the subject’s renal disease. To identify the most probable disease-causing mutation, we used the following criteria for exome variant filtering^40–43^: (1) exclusion of all variants that did not change the amino-acid sequence or affected canonical splice sites (defined as ± 6 nucleotides surrounding the exon-intron boundary), (2) exclusion of variants reported in the homozygous state or with a minor allele frequency greater than 0.1% in a control cohort (ExAC and gnomAD genome databases), (3) inclusion of homozygous bi-allelic variants with appropriate parental segregation consistent with our above hypothesis, and (4) assessment of variants for deleteriousness based on *in silico* prediction of their impact on protein structure and/or splice site function. This approach, however, failed to yield a strong candidate mutation. Therefore, trio whole genome sequencing (TRIO-WGS) was performed in order to extend coverage to non-coding genome regions.

**Figure 5.**
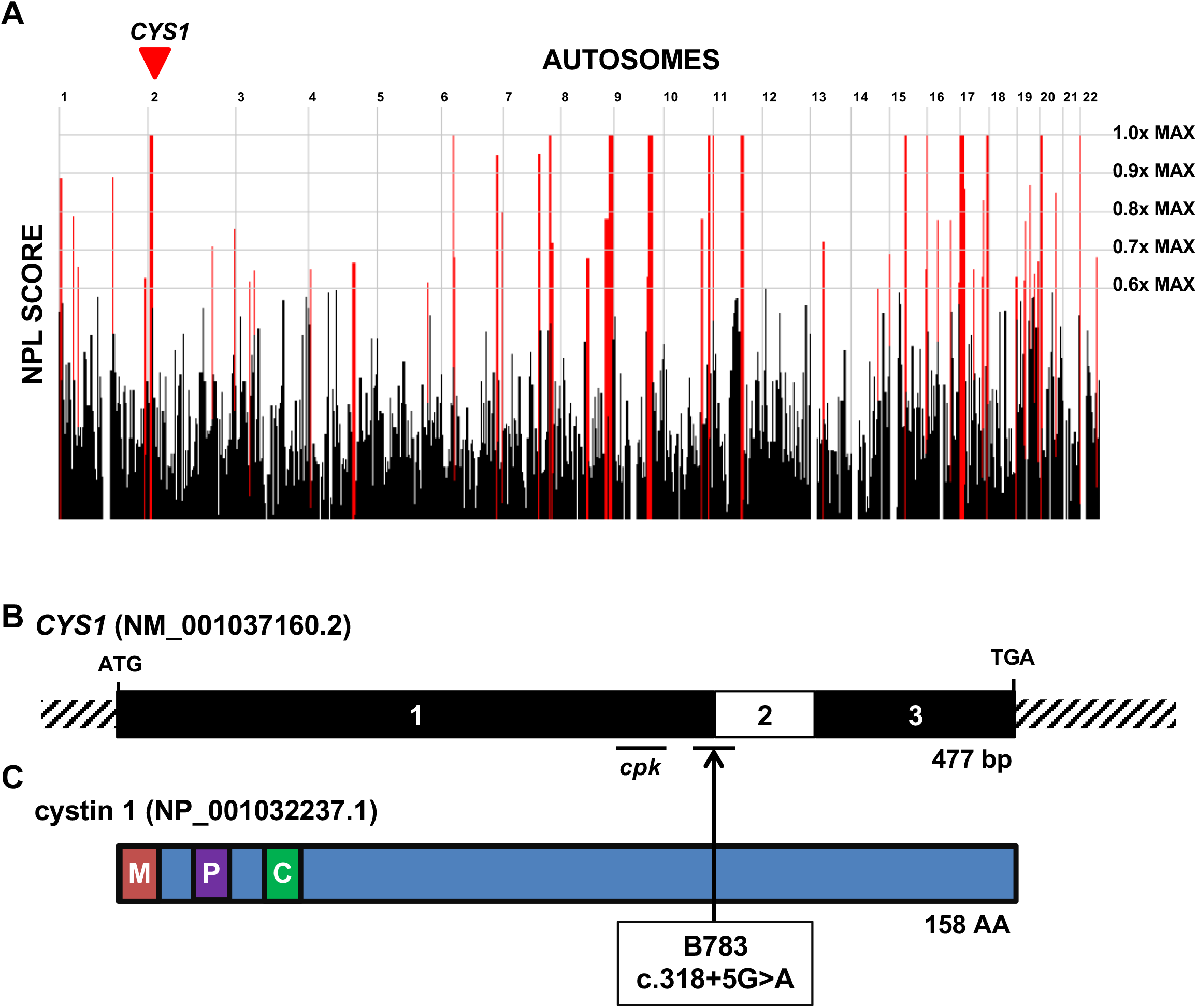
Whole genome sequencing identified a recessive mutation in the gene *CYS1* in one family with polycystic kidneys and liver fibrosis. (**A**) Genome-wide homozygosity mapping in individual B783 identifies homozygous peak regions. The gene locus for *CYS1* is located within a peak region of chromosome 2 (arrowhead). (**B**) Exon structure of *CYS1* cDNA. Positions of start codon, stop codons, and of the affected splice site (boundary exon 1-2) are indicated. The region within exon 1 deleted in the *cpk* mouse is also indicated. (**C**) Domain structure of the encoded cystin-1 protein, which contains a myristoylation site (M), polybasic region (P) and AxEGG motif required for targeting cystin-1 to the cilium (C). The arrow indicates the boundary between protein regions encoded by exons 1 and 2. Abbreviations: AA, amino acid; BP, base pairs; C, cilium trafficking domain; M, myristoylation site; P, polybasic region.

Employing the same filtering criteria described for TRIO-WES led to the identification of a mutation lying in a region of homozygosity of descent (Figure 5A) representing a splice-site mutation (c.318+5G>A) in the exon 1 donor site of *CYS1* (Figure 5B,C). The following criteria strongly suggest a deleterious mutation: (i) the splice variant is extremely rare, being absent from the gnomAD database (gnomAD, version 2.1.1), and (ii) four independent *in silico* prediction tools indicate a deleterious effect of the mutation on splicing (Table 1). Importantly, neither TRIO-WES nor TRIO-WGS analysis revealed causative mutations in >100 cystic kidney disease genes, including *PKHD1* and *DZIP1L*. Allele-specific PCR analysis confirmed homozygosity and heterozygosity of the mutation in the proband and both parents, respectively (Supplementary Figure S1).

**Table 1.** *CYS1* mutation in one family with an ARPKD phenotype.

| Family | Change | Affect | Exon<br>(Zyg, Seg) | *In silico*<br>Severity<br>Scores | gnomAD<br>(H/h/T) | Sex | Ethnic<br>origin | PC | Clinical Phenotype |
| --- | --- | --- | --- | --- | --- | --- | --- | --- | --- |
| B783 | c.318+5<br>G>A | Splice | 1<br>(Hom, B) | CADD<br>17.06<sup>a</sup><br>HSF -13.17<sup>bc</sup><br>MaxEnt -<br>62.8%<sup>c</sup><br>NNS -71.3%<sup>c</sup> | 0/0/<br>~251,000 | M | Egypt | Y | <u>Initial Onset</u>: 5 years,<br>polyuria, polydipsia<br><u>Serum Studies</u>: Cr 0.6<br>mg/dL<br><u>RUS</u>: bilateral cortical and<br>medullary cysts<br><u>Extra-renal</u>: Mild congenital<br>hepatic fibrosis |

Abbreviations: B, both parents appropriately heterozygous; CADD, Combined Annotation-Dependent Depletion prediction score; Cr, serum creatinine; gnomAD, Genome Aggregation database; H, homozygotes in gnomAD; h, heterozygous alleles in gnomAD; Hom, homozygous zygosity; HSF, HSF splice prediction score; M, male; MaxEnt, MaxEnt splice prediction score; NNS, NNSPLICE splice-site mutation prediction score; PC, parental consanguinity; RUS, renal ultrasound; Seg, segregation; T, total alleles in gnomAD; Y, yes; Zyg, zygosity.
<sup>a</sup>CADD scores between 10-20 are in the 1-10% most deleterious substitutions possible in human genome in terms of predicted effect on the gene product.
<sup>b</sup>Reports wildtype site broken by this change.
<sup>c</sup>Percentage reflects expected reduction in splicing at this donor site as consequence of change.

In light of the above genetic findings, a further evaluation of the clinical history of subject B783 revealed mild congenital liver fibrosis, which is also consistent with the established phenotype of the *cpk* mouse.^44–46^ Our findings strongly support the first identification of a causative mutation in *CYS1* in a human patient with ARPKD.

## Discussion

In the current study, we demonstrate that expression of the *Cys1* transgene in renal collecting ducts of *cpk* mice rescues the cystic phenotype and down-regulates expression of *Myc in vivo*. We also report the first ARPKD patient with a homozygous *CYS1* mutation, a c.818+5G>A variant predicted to disrupt splicing. This variant affects the +5-position of the canonical donor splice site, AG/GURAGU. Pathogenic variants affecting +5 position of the donor splice site have been reported in other genes^47–49^ and detailed analysis of G-to-A sequence changes at the +5 position have revealed disrupted base pairing between donor splice site of a pre-mRNA and the U1snRNP of the spliceosome leading to decreased efficiency of the splice site recognition and exon skipping.^50^

The identification of a pathogenic *CYS1* variant in a patient with an ARPKD-like phenotype confirms the importance of the *CYS1* gene product for normal function of human collecting duct cells. While it is surprising that *CYS1* deficiency causing ARPKD has been observed in only one family, it is important to note that this gene is GC-rich, particularly its first exon (Figure S3). Such GC-rich regions can be difficult to amplify and sequence using Sanger methodology^51^ and can be missed in next-generation sequencing because low sequence complexity prevents efficient capture prior to library construction.^52^

Development of collecting duct cysts in humans and mice with cystin deficiency suggests a shared pathobiology and possibly similar molecular mechanisms underlying cyst formation. In the *cpk* mouse model, renal cysts initially develop at embryonic day 15.5 (E15.5) and are restricted to proximal tubules.^53,54^ As *cpk* mice develop and disease progresses, cysts predominantly affect the distal collecting duct region.^53^ Specific expression of cystin-GFP in developing ureteric bud-derived collecting ducts rescued the renal cystic phenotype. Transgenic *cpk* mice expressing the fusion protein only in AQP2-positive collecting duct cells exhibited survival rates and kidney sizes similar to wild type mice. Interestingly, while collecting duct cysts were absent in rescued *cpk* mice, these animals did express proximal tubular cysts, suggesting that the initial phase of proximal tubular cystogenesis was not rescued. Proximal tubule cysts have also been observed in human ARPKD fetal specimens between 14 and 26 weeks of gestation, but not in the kidneys of fetuses older than 34 weeks of gestation.^39^ These observations suggest a gradual shift of cyst formation sites from proximal tubules to collecting ducts during the early fetal development in both human and mouse ARPKD.

Cystin is a cilium-associated protein that localizes to the basal bodies and the ciliary axoneme.^17,18,55^ Treatment of *cpk* mice with paclitaxel, which promotes microtubule assembly, prevents renal cyst formation, suggesting that cystin may stabilize microtubule assembly within the ciliary axoneme.^56^ The primary cilia of the collecting duct epithelium function as transmitters of mechano- and chemosensory stimuli to signaling pathways that regulate multiple key cellular processes including differentiation, proliferation, apoptosis, tissue homeostasis and cell polarity.^57^ We have previously demonstrated that cystin, with two functional nuclear localization signals, can be released from the ciliary membrane through a myristoyl-electrostatic switch and translocate to the nucleus where it forms a regulatory complex with necdin to modulate *Myc* expression.^24^

The *Myc* proto-oncogene plays a critical role in normal kidney development^58^ and several lines of evidence suggest a central role for dysregulated *Myc* expression in the pathophysiology of polycystic kidney disease. First, overexpression of *Myc* in the kidneys of SBM transgenic mice causes polycystic kidney disease.^59^ Furthermore, renal cystic disease remitted in SBM mice that underwent spontaneous reversion to normal kidney *Myc* expression.^60^ Second, treatment of *cpk* mice with antisense *Myc* oligonucleotides mitigated the cystic phenotype.^22^ Third, *Myc* is overexpressed in mouse models of autosomal dominant polycystic kidney disease (ADPKD)^61,62^ and *Myc* expression appears to be tightly regulated by PC1, the product of the *Pkd1* gene.^63^ Fourth, pharmacological inhibition of glucogen synthase kinase 3 beta (GSK3beta), which accelerates cyst formation in *cpk* mice, leads to decreased *My*c expression and amelioration of the cystic phenotype.^64^ Similarly, *Myc* is down-regulated in *Cys1*^*cpk*^/*cpk;Smad3*^+/−^ mice and such double mutants have a milder phenotype than *cpk* mice.^25^ Our findings that complementation of mutant *cpk* with cystin-GFP rescues the cystic phenotype and restores normal *Myc* expression confirms that cystin acts *in vivo* as a negative regulator of *Myc*.

In summary, we demonstrate that cystin deficiency causes an ARPKD-like phenotype in mice that can be rescued by targeted renal expression of a cystin-GFP fusion protein, most probably by downregulating *Myc* expression in collecting duct cells. Our identification of the first case of human ARPKD associated with a *CYS1* mutation confirms the relevance of the *cpk* mouse as an ARPKD model yielding important insights into molecular mechanisms underlying disease pathobiology.

## Methods

### Animal study approvals

All mouse experiments were approved by the Institutional Animal Care and Use Committees of Children’s National Research Institute and the University of Alabama at Birmingham (UAB). Knock-in transgenic mice were generated at the University of Alabama at Birmingham (UAB) Transgenic & Genetically Engineered Models Core facility. *Ksp-Cre* mice were obtained from Jackson Laboratory (Bar Harbor, ME). Mouse colonies were maintained in the animal facility at Children’s National Research Institute.

### Antibodies and lectins

Anti-GAPDH antibody was purchased from Cell Signaling Technologies (# 2118). Anti-AQP2 antibody was purchased from Santa Cruz Biotechnologies (# SC9882). Alexa Fluor 488 conjugated anti-GFP antibody was obtained from Life Technologies (# A21311). Polyclonal rabbit anti-cystin antibody (70053) was generated in our lab and described previously.^18^ Rabbit monoclonal anti-c-Myc antibody was purchased from Abcam (# ab32072). Goat anti-rabbit HRP conjugated secondary antibody was purchased from American Qualex Solution Products (# A102PS). Donkey anti-Goat IgG Alexa Fluor 555 was obtained from Life Technologies (# A21432). Lectins LTA-FITC (# W0909) and DBA-Rhodamine (# Y0828) were obtained from Vector Laboratories.

### Vector cloning

*Cys1-GFP* cDNA was amplified from previously described pEGFP-N1.^18^ Gateway PCR primers were used to add flanking attB sites. A one-tube Gateway reaction was performed using pDonr221 and pRosa26 Dest.^65^ The reaction product was used to transform competent STBL3 cells (Life Technologies, # C7373-03) that were plated on Ampicillin and Kanamycin to select for destination and entry clones, respectively. Destination clones were screened by restriction enzyme digest prior to sequencing. The pRosa26 Dest *Cys1-GFP* targeting vector was linearized with *Kpn1* and electroporated into ES cells. G418 resistant colonies were screened by long-range PCR as described^66^ and positive clones were used to produce chimeric founder mice.

### PCR genotyping

PCR conditions are described in Supplemental Table 1.

### Immunoblotting

Kidney tissue was collected, homogenized, and processed for immunoblotting as previously described.^24^ For cystin and control western blots, immuno-reactive protein bands were visualized using SuperSignal West Dura chemiluminescent substrate (Thermo Fisher Scientific, # 34076) and exposed to film. For C-MYC and control western blots, images were obtained with ChemiDoc Imaging System (Bio-Rad laboratory, Inc.) Densitometry was analyzed using Image Lab (Bio-Rad laboratory, Inc., Version 6.0).

### Kidney histology

Tissue samples were collected and fixed in 10% formalin (Fisher Scientific, # 23-245-684) for 2 days, then stored in 70% ethanol. The samples were dehydrated, paraffin embedded, cut into 5 μm sections and stained with hematoxylin and eosin (H&E) and slide-mounted for examination by the UAB Comparative Pathology Laboratory.

### Immunofluorescence analysis

Tissue samples were collected from 6 week-old mice and processed, using published methods.^33^ Immunofluorescence detection and image acquisition were performed using an Olympus FLUOVIEW FV1000 confocal laser scanning microscope configured with both an Argon Laser (488 nm) and a Laser diode (405 nm, 440 nm, and 559 nm). Images were analyzed using Olympus FV10-ASW 3.0 Viewer software.

### Lectin staining

Five μm sections of fixed paraffin embedded kidney tissues (prepared as described for histopathology) were stained with Rhodamine labeled DBA (Vector Laboratory, # RL-1032) and Fluorescein labeled LTA (Vector Laboratory, # FL-1321).^67^ Immunofluorescence detection, image acquisition, and analysis were performed as described above.

### Human study approval

Approval for human subjects research was obtained from Institutional Review Boards of the University of Michigan, Boston Children’s Hospital, and local IRB equivalents.

### Human research subjects

Blood samples and pedigrees were obtained following informed consent from individuals with cystic kidney disease or their legal guardians. The diagnosis of cystic kidney disease was based on published clinical criteria. Clinical data were obtained using a standardized questionnaire (http://www.renalgenes.org).

### Whole genome sequencing and mutation calling

TRIO-WES and data processing were performed by the Genomics Platform at the Broad Institute of Harvard and MIT (Broad Institute, Cambridge, MA). Exome sequencing (>250 ng of DNA, at >2 ng/μl) was performed using Illumina exome capture (38 Mb target). Single nucleotide polymorphisms (SNPs) and insertions/deletions (indels) were jointly called across all samples using the Genome Analysis Toolkit (GATK) HaplotypeCaller. Default filters were applied to SNP and indel calls using the GATK Variant Quality Score Recalibration approach. Lastly, variants were annotated using the Variant Effect Predictor. For additional information, refer to the Supporting Information Section S1 in the exome aggregation consortium (ExAC) study.^68^ The variant call set was uploaded on to Seqr (https://seqr.broadinstitute.org) and analysis of the entire WES output was performed. TRIO-WGS and data processing were performed by the Genomics Platform at the Broad Institute of MIT and Harvard. PCR-free preparation of sample DNA (350 ng input at >2 ng/ul) was accomplished using Illumina HiSeq X Ten v2 chemistry. Libraries were sequenced to a mean target coverage of >30x. Genome sequencing data was processed through a pipeline based on Picard, using base quality score recalibration and local realignment at known indels. The BWA aligner was used for mapping reads to the human genome build 38. Single Nucleotide Variants (SNVs) and insertions/deletions (indels) were jointly called across all samples using Genome Analysis Toolkit (GATK) HaplotypeCaller package version 3.4. Default filters were applied to SNV and indel calls using the GATK Variant Quality Score Recalibration (VQSR) approach. Annotation was performed using Variant Effect Predictor (VEP). Lastly, the variant call set was uploaded to seqr for collaborative analysis between the CMG and investigator.

Mutation calling was performed in line with proposed guidelines,^41^ and the following criteria were employed as previously described^42,43^. The variants included were rare in the population with mean allele frequency <0.1% and with 0 homozygotes in the adult reference genome databases ExAC and gnomAD. Additionally, variants were non-synonymous and/or located within splice-sites. Based on an autosomal homozygous recessive hypothesis, homozygous variants were evaluated. Subsequently, variant severity was classified based on prediction of protein impact (truncating frameshift or nonsense mutations, essential or extended splice-site mutations, and missense mutations). Splice-site mutations were assessed by *in silico* tools MaxEnt, NNSPLICE, HSF, and CADD splice-site mutation prediction scores.^69–72^ Missense mutations were assessed based on SIFT, MutationTaster and PolyPhen 2.0 conservation prediction scores^73–75^ and evolutionary conservation based on manually derived multiple sequence alignments.

### Homozygosity mapping (HM)

Homozygosity mapping was calculated based on whole exome sequencing data. In brief, aligned BAM files were processed using Picard and SAMtools4 as described previously.^76^ Single nucleotide variant calling was performed using Genome Analysis Tool Kit (GATK).^77^ The resulting VCF files were used to generate homozygosity mapping data and visual outputs using the program Homozygosity Mapper.^78^

### Web resources

UCSC Genome Browser, genome.ucsc.edu

Ensembl Genome Browser, www.ensembl.org

gnomAD browser 2.0.3., gnomad.broadinstitute.org

Polyphen2, genetics.bwh.harvard.edu/pph2

Sorting Intolerant From Tolerant (SIFT), sift.jcvi.org

MutationTaster, www.mutationtaster.org

Combined Annotation Dependent Depletion, cadd.gs.washington.edu

NNSPLICE splice-site mutation prediction, www.fruitfly.org/seq_tools/splice.html

MaxEnt splice prediction, hollywood.mit.edu/burgelab/maxent/Xmaxentscan_scoreseq_acc.html

Human Splice Finder, www.umd.be/HSF/

## Acknowledgements

The authors thank Gene Siegel, MD, PhD (UAB) for histological analysis and the Cellular Imaging and Analysis Core at Children’s National Research Institute for assistance with microscopy. This work was supported by National Institute of Diabetes and Digestive and Kidney Diseases (NIDDK) grant P30 DK074038. The authors would like to thank members of the University of Alabama at Birmingham Transgenic & Genetically Engineered Models (TGEMs) facility for creating the Cys1-GFP transgene knock-in animals. TGEMs is supported by NIH National Cancer Institute Grant P30CA13148, NIH NIAMS Grant P30AR048311, and NIH NIDDK Grants P30 DK074038, P30 DK05336, and P60 DK079626 (to RAK). F.H. is the William E. Harmon Professor of Pediatrics. This research is supported by a grant from the National Institutes of Health to F.H. (DK-076683-13). A.J.M. was supported by an NIH Training Grant (T32DK-007726), by the 2017 Post-doctoral Fellowship Grant from the Harvard Stem Cell Institute, and by the American Society of Nephrology Lipps Research Program 2018 Polycystic Kidney Disease Foundation Jared J. Grantham Research Fellowship. F.B. was supported by a fellowship grant (404527522) from the German Research Foundation (DFG). Sequencing and analysis were provided by the Broad Institute of MIT and Harvard Center for Mendelian Genomics (Broad CMG) and was funded by the National Human Genome Research Institute, the National Eye Institute, and the National Heart, Lung and Blood Institute grant UM1 HG008900 and in part by National Human Genome Research Institute grant R01 HG009141.

---

**Supplementary Figure S1:**
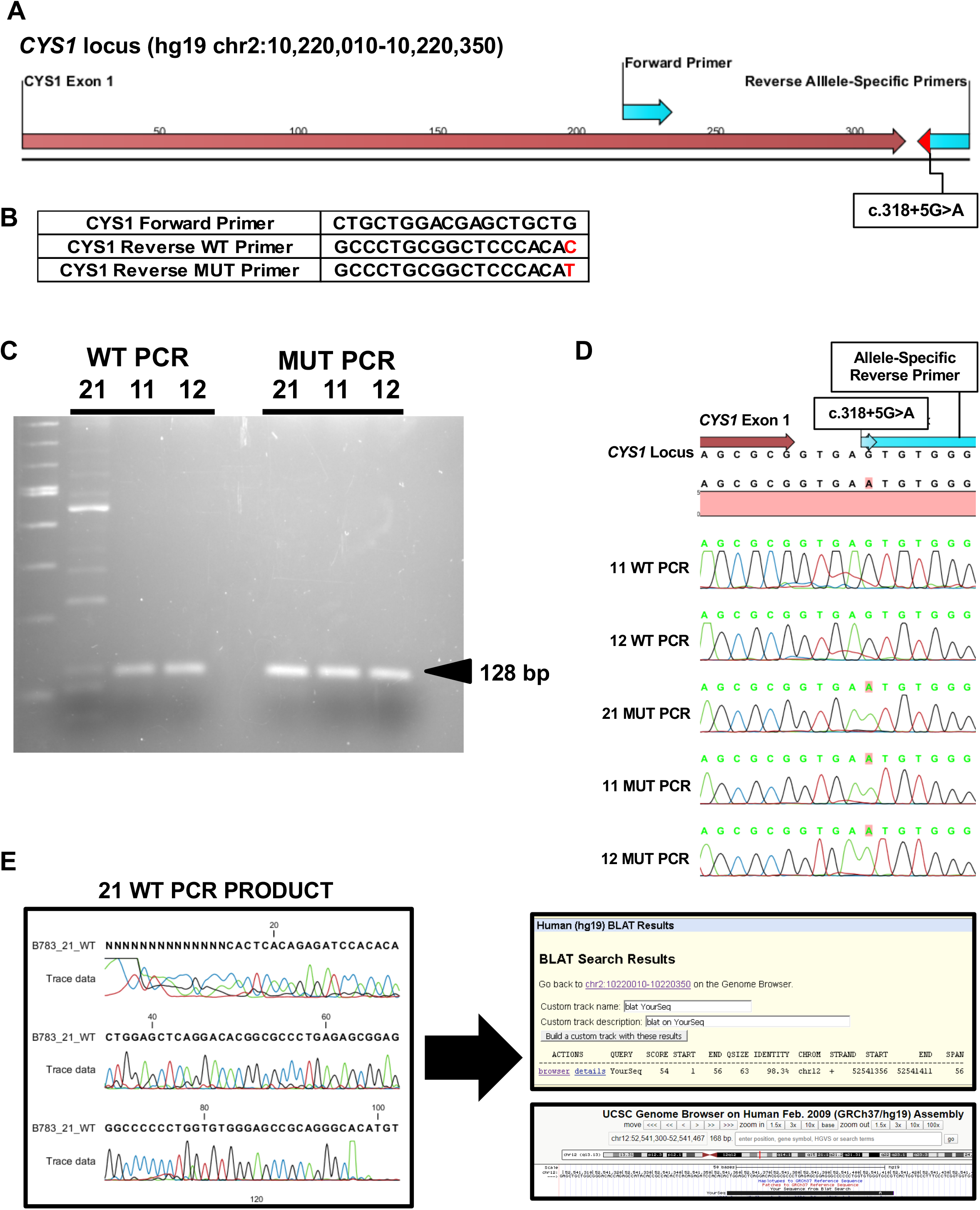
Confirmation of *CYS1* mutation in family B783 with childhood onset polycystic kidney disease and liver fibrosis. (**A**) Map showing primer location in context of *CYS1* variant for allele-specific primer PCR. (**B**) Table shows shared forward primer and allele specific reverse primers for the wildtype (WT) and mutated nucleotide (MUT). (**C**) Gel electrophoresis of *CYS1* allele-specific PCR for WT and MUT alleles from DNA of the proband (21), father (11), and mother (12) in family B783 is shown. PCR products were noted at the expected size (128 bp) for both parents for the WT PCR, while the WT PCR in the affected child yielded only non-specific bands. The MUT PCR yielded a single product of expected size in all three family members. This suggested the parents were heterozygous for the MUT allele, while the affected proband is homozygous. (**D**) The PCR products were excised at 128 bp for each of the 6 reactions in (C) and were sequenced using the Sanger approach. The sequences derived from the chromatograms were aligned to the *CYS1* locus as shown. The WT and MUT products from parental DNA (11 and 12) and the MUT product for the affected proband (21) confirmed as amplification products of this locus. The product from WT PCR of the affected proband (21) DNA did not align. (**E**) The chromatogram sequence from the WT PCR for the proband was aligned to the human genome using UCSC Browser BLAT. This yielded only one search result, indicating 98.3% similarity with an intron in chromosome 12. This supports that the WT PCR from the DNA of the affected proband yielded non-specific products, while the others yielded only the *CYS1* locus. Abbreviations: 11, father; 12, mother, 21, proband; WT, wildtype; MUT, mutant.

